# Chromosome-scale genome assembly of *Poa trivialis* and population genomics reveals widespread gene flow in a cool-season grass seed production system

**DOI:** 10.1101/2023.11.03.565483

**Authors:** Caio A. C. G. Brunharo, Christopher W. Benson, David R. Huff, Jesse R. Lasky

## Abstract

*Poa trivialis* (L.) is a cool-season grass species found in various environments worldwide. In addition to being a desired turfgrass species, it is a common weed of agricultural systems and natural areas. As a weed, it is an important contaminant of commercial cool-season grass seed lots, resulting in widespread gene flow facilitated by human activities and causing significant economic losses to farmers. To better understand and manage infestations, we assembled and annotated a haploid genome of *P. trivialis* and studied troublesome field populations from Oregon, the largest cool-season grass seed producing region in the United States. The genome assembly resulted in 1.35 Gb of DNA sequence distributed among seven chromosome-scale scaffolds, revealing a high content of transposable elements, conserved synteny with *P. annua*, and a close relationship with other C_3_ grasses. A reduced-representation sequencing analysis of field populations revealed limited genetic diversity and suggested potential gene flow and human-assisted dispersal in the region. The genetic resources and insights into *P. trivialis* provided by this study will improve weed management strategies and enable the development of molecular detection tests for contaminated seed lots to limit seed-mediated gene flow. These resources should also be beneficial for turfgrass breeders seeking to improve desirable traits of commercial *P. trivialis* varieties and help to guide breeding efforts in other crops to enhance the resiliency of agricultural ecosystems under climate change.

**Significance statement:** The chromosome-scale assembly of *Poa trivialis* and population genomic analyses provide crucial insights into the gene flow of weedy populations and contribute a valuable genomic resource for the plant science community.

## Introduction

*Poa trivialis* (L.) is a cool-season grass species that has successfully colonized a diverse range of environments. Native to Eurasia, it can be found in all continents except Antarctic. *Poa trivialis* is considered a weed of agricultural crops (George, 1990). Because it can reproduce by both seed and stolons, it is often difficult to control with conventional management practices, such as herbicide applications (Haggar, 1971).

Weeds pose a significant threat to food security and the sustainability of agroecosystems. Over the last century, turfgrass seed, particularly Kentucky bluegrass (*P. pratensis*) has been intentionally imported from Europe and grown around the world for various purposes. Being a closely related species with similar growth patterns and seed shape, *P. trivialis* seed is a common contaminant of Kentucky bluegrass seed lots, resulting in the broad dispersal of *P. trivialis*. Undesired infestations of *P. trivialis* in fields can directly and indirectly cause damage to desired vegetation. As such, *P. trivialis* can be particularly problematic when infesting cool-season grass seed crops, such as perennial ryegrass (*Lolium perenne*), annual ryegrass (*L. multiflorum*), tall fescue (*Schenodorum arundinacea*), and Kentucky bluegrass and can cause concerns with contamination of commercial seed lots (Haggar, 1979).

In addition to the undesirable spread of *P. trivialis* as a seed contaminant, this species has been broadly introduced intentionally with uses in agriculture as permanent pasture as well as a turfgrass species. *Poa trivialis* is adapted to moist, shaded areas, with relatively limited tolerance to drought. In the southern United States, this species has been utilized as an overseeding species for dormant warm-season turfgrasses like bermudagrass (*Cynodon spp.*) on golf courses. In these turfgrass systems, managers sow a cool-season (i.e., winter-adapted) species, such as *P. trivialis,* in the fall when temperatures start to decrease, just as the warm-season (i.e., summer-adapted) turfgrass species start to enter winter dormancy. The shallow root system and limited tolerance to high temperatures of *P. trivialis* allows the warm-season turfgrass to outcompete and resume growth in the spring (Hurley, 2003).

Over the last few years, practitioners have reported an increase in *P. trivialis* infestation in the cool-season grass seed crops in Oregon, one of the largest cool-season grass seed producing regions in the world. In grass seed crops, farmers must adhere to stringent seed purity regulations to certify their seed lots and maintain a profitable production system. Grass seed crops from Oregon are sold as planting seed to establish turf throughout the USA and the world for home lawns, golf courses, and other sports fields. Commercial seed lots that are contaminated with *P. trivialis* can cause lost revenue and be a source for long-distance weed dispersal. To illustrate the potential of seed contamination, a survey was conducted by Levy (1998) on commercial seed lots of creeping bentgrass (*Agrostis stolonifera*) and found contamination with *P. trivialis* in 30% of samples.

Limited genetic information is available on the biology of *P. trivialis*, and most research to date has focused on this species as a crop rather than as a weed. Ahmed et al. (1972) identified 14 chromosomes in *P. trivialis* (2n = 2x = 14), where most seem to have similar sizes, and their data indicated that this species in unlikely to be a progenitor to the closely related tetraploid, *P. annua*. In fact, Mao & Huff (2012) suggested that *P. trivialis* is phylogenetically closer related to *P. hybrida* than to *P. annua* or its progenitors, *P. infirma* and *P. supina*. In studying 27 populations of a global germplasm of cultivated and wild accessions of *P. trivialis*, Rajasekar et al. (2006) found a high degree of genetic diversity. The authors observed that the within-accession genetic variation accounted for most of the total genetic variation (>87%), aligning with expectations based on its self-incompatible and obligate outcrossing breeding system. They also observed a wide range of between-accession genetic diversity depending on the comparison. For instance, an accession from the USA was closely related to an accession from Iraq, whereas this same accession had a much greater similarity coefficient compared to another population from the USA, suggesting that intercontinental gene flow likely shaped the tested populations.

Generating genomic resources for *P. trivialis* will help researchers to better inform management practices for farmers, improve this species as a desirable turfgrass, and understand the underlying biology that has allowed it to colonize a wide variety of environments. Furthermore, examining the genetic diversity within and between populations of *P. trivialis* is essential to understanding patterns of dispersal, evolution of weedy traits, and ultimately to develop molecular detection tests to limit contamination and gene flow. In this context, the objective of this study was to further our understanding of the genetics of weedy *P. trivialis* from Oregon. Because of the limited genetic resources available for *P. trivialis*, we first assembled a reference genome and conducted a reduced-representation sequencing experiment to elucidate the genetic relationship within and between 12 *P. trivialis* populations collected from agricultural fields.

## Results

### Reference genome assembly and annotation

The first step towards improving genomic resources for *P. trivialis* was to assemble a haploid chromosome-level reference genome. This was achieved by generating approximately 68 Gb of highly accurate (>99%) PacBio HiFi long-read data (equivalent to 25× coverage), and 114 Gb of Illumina paired-end sequencing data from a proximity ligation library (Hi-C). The genome was assembled with *hifiasm* 0.16.1-r375 (Cheng et al., 2021) in integrated mode using both HiFi long-read and Hi-C data. The genome assembly was 1.39 Gb in size across 846 contigs, of which 97% of the DNA content was placed in 7 scaffolds (L_95_ = 7), indicating our approach resulted in a reference quality, chromosome-scale assembly (Table 1, Figure 1). The assembly size was cross-validated using *k*-mer analysis and flow cytometry, where both suggested a haploid genome size of 1.35 Gb. In addition, *k*-mer analysis indicated a 2.53% heterozygosity. We performed a BUSCO (Manni et al., 2021) analysis on the assembly to identify single-copy orthologs in the genome and used the *Poales* lineage for comparison. This analysis indicated that >95% of conserved *Poales* orthologs were represented as single copies in the *P. trivialis* reference genome. The remaining BUSCOs were either duplicated (4.4%) or fragmented and missing (1%) (Table 1).

**Figure 1.**
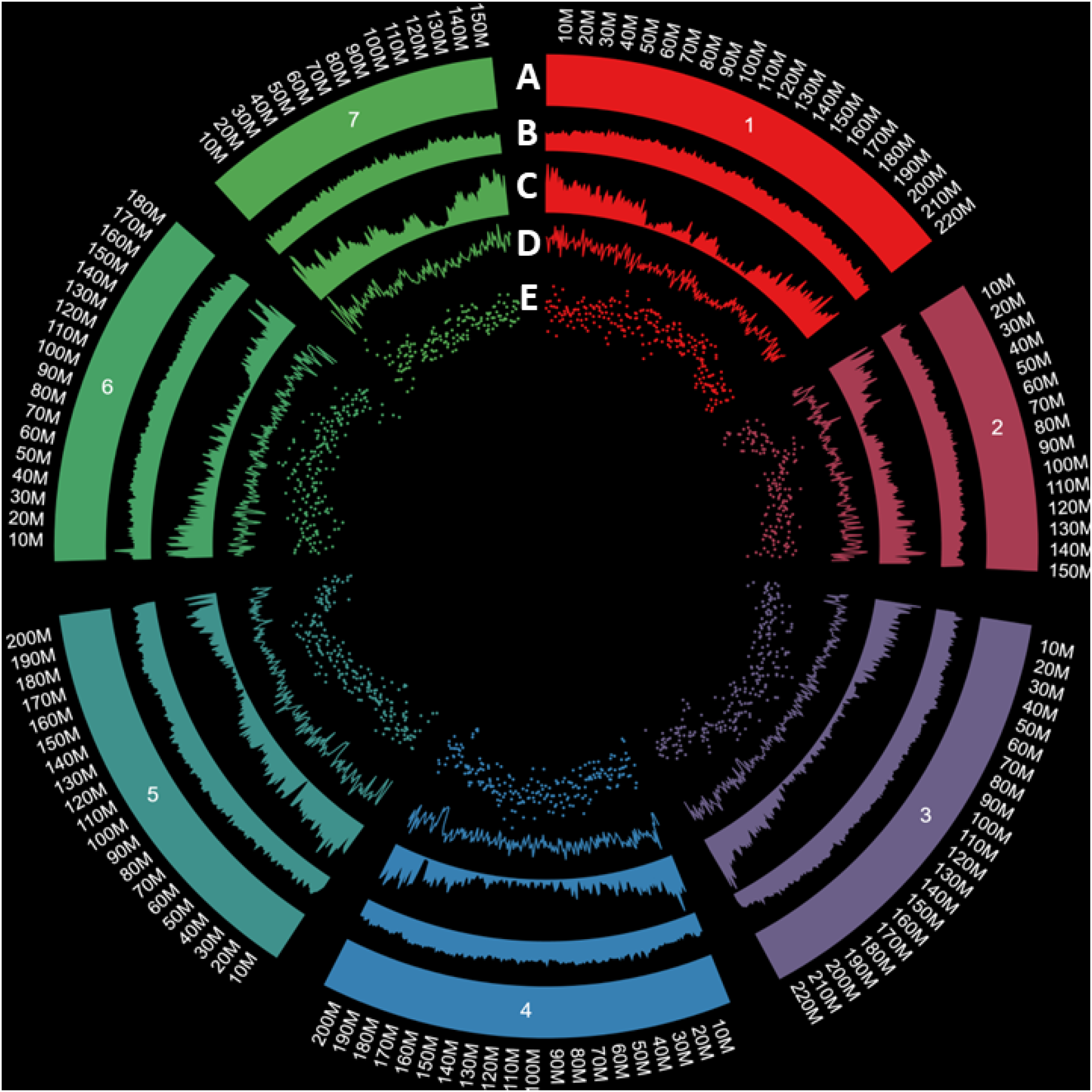
Overview of the *P. trivialis* genome with a *circos* graph. (A) Seven chromosomes assembled, (B) distribution of repetitive elements, (C) location of the gene content, (D) distribution of small insertions and deletions, and (E) single-nucleotide polymorphisms. Outer numbers represent physical location in mega bases.

**Table 1.** Summary statistics of *P. trivialis* genome assembly.

| Assembly parameter | Estimates |
| --- | --- |
| # contigs | 846 |
| # contigs ($\geq 25,000$ bp) | 741 |
| Largest contig | 225,343,272 |
| Total length | 1,397,906,150 |
| Total length ($\geq 25,000$ bp) | 1,395,716,373 |
| N<sub>50</sub> | 209,324,251 |
| N<sub>90</sub> | 150,158,976 |
| auN | 192,269,589 |
| L<sub>50</sub> | 4 |
| L<sub>95</sub> | 7 |
| GC (%) | 44.18 |
| BUSCO |  |
| Complete and single-copy (%) | 95.6 |
| Complete and Duplicated (%) | 4.4 |
| Fragmented (%) | 0.5 |
| Missing (%) | 0.03 |
| Number of genes identified | 58,464 |

To annotate the *P. trivialis* genome, we used several datasets. First, we sequenced two IsoSeq libraries using RNA extracted from leaf and root tissues using the PacBio Sequel IIe platform. The *isoseq3* (https://github.com/PacificBiosciences/IsoSeq) and *TAMA* (Kuo et al., 2020) pipelines were implemented to process highly accurate full-length transcripts (nearly 2.1 M high-quality reads) into a transcriptome with reduced redundancy. Second, a non-redundant transposable element library was created by providing the *P. trivialis* reference genome and transcriptome as inputs into *Extensive De-novo TE Annotator* (EDTA) (Ou et al., 2019), with supplemental element identification with RepeatModeler (Flynn et al., 2020). This step suggested that >77% of the *P. trivialis* genome is composed of transposable elements, with the majority (39%) being Gypsy-type retrotransposons (Table 2). Third, gene models were created with SNAP (Korf, 2004) and Augustus (Stanke et al., 2006). Finally, these datasets, along with the proteomes of *Brachypodium distachyon*, wheat (*Triticum aestivum*), and barley (*Hordeum vulgare*) were used to perform a homology-based gene prediction with MAKER (Campbell et al., 2014). Our pipeline resulted in the annotation of 58k genes with an average length of 2.7 kb.

**Table 2.** The landscape of transposable elements in *P. trivialis*.

| Class | Count | bp masked | %masked |
| --- | --- | --- | --- |
| LTR |  |  |  |
| Copia | 113729 | 100,178,374 | 7.17 |
| Gypsy | 582547 | 548,647,164 | 39.25 |
| unknown | 432532 | 186,617,080 | 13.35 |
| TIR |  |  |  |
| CACTA | 101042 | 56,700,805 | 4.06 |
| Mutator | 82410 | 34,627,136 | 2.48 |
| PIF_Harbinger | 31951 | 15,665,832 | 1.12 |
| Tc1_Mariner | 47430 | 12,955,630 | 0.93 |
| hAT | 24366 | 8,544,389 | 0.61 |
| nonLTR |  |  |  |
| LINE_element | 4240 | 2,205,638 | 0.16 |
| unknown | 309 | 376,291 | 0.03 |
| nonTIR |  |  |  |
| helitron | 127847 | 69,971,256 | 5.01 |
| Repeat region | 111610 | 44,843,398 | 3.21 |
| Total | 1660069 | 1,081,353,823 | 77.36 |

### Comparative genomics between P. trivialis and species of agricultural interest

The genomes of species of agronomic interest and potentially close relatives to *P. trivialis* were compared. First, we analyzed the genomes of *P. trivialis*, *P. infirma, B. distachyon,* and rice (*Oryza sativa*) with CoGe’s SynMap tool (Haug-Baltzell et al., 2017). We calculated the synonymous substitution rates (K_s_) between homologous gene pairs and, based on a mutational rate of 5.76174 × 10^-9^ substitutions per site per year (De La Torre et al., 2017) and estimated that *P. trivialis* diverged from *P. infirma*, *B. distachyon* and *O. sativa* 12.4, 28.7, and 46.2 M years ago, respectively. Pairs of colinear and intra-genomic genes in the *P. trivialis* genome revealed dispersed K_s_, suggesting that a species-specific whole-genome duplication did not occur in this grass. A phylogenetic analysis of *P. trivialis* was performed with a larger number of grass species and further elucidated their evolutionary relationship (Figure 2). A synteny analysis between the genomes of diploid *P. trivialis* (1.35 Gb) and tetraploid *P. annua* (1.78 Gb) revealed that both species have conserved gene orders along most chromosomes in a predictable 1:2 relationship. Synteny analysis also identify several chromosomal rearrangements. For example, a small inversion was observed between *P. trivialis*’ chr1 and *P. annua*’s 1A, chr2 and 2B, chr4 and 4A and 4B, and a large inversion between chr5 and 5B. A large translocation was observed between chr7 and 2A and 2B (Figure 2C).

**Figure 2.**
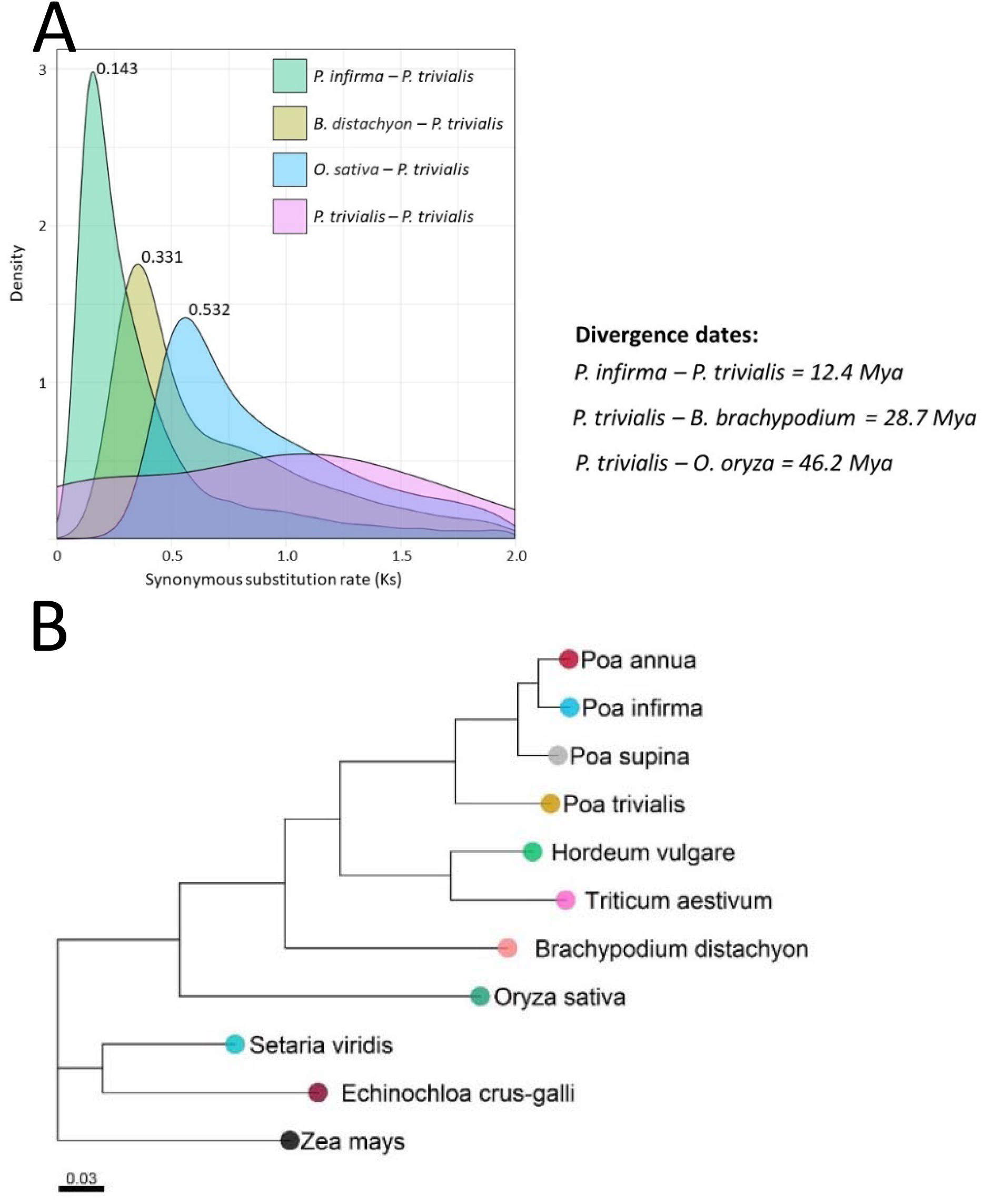
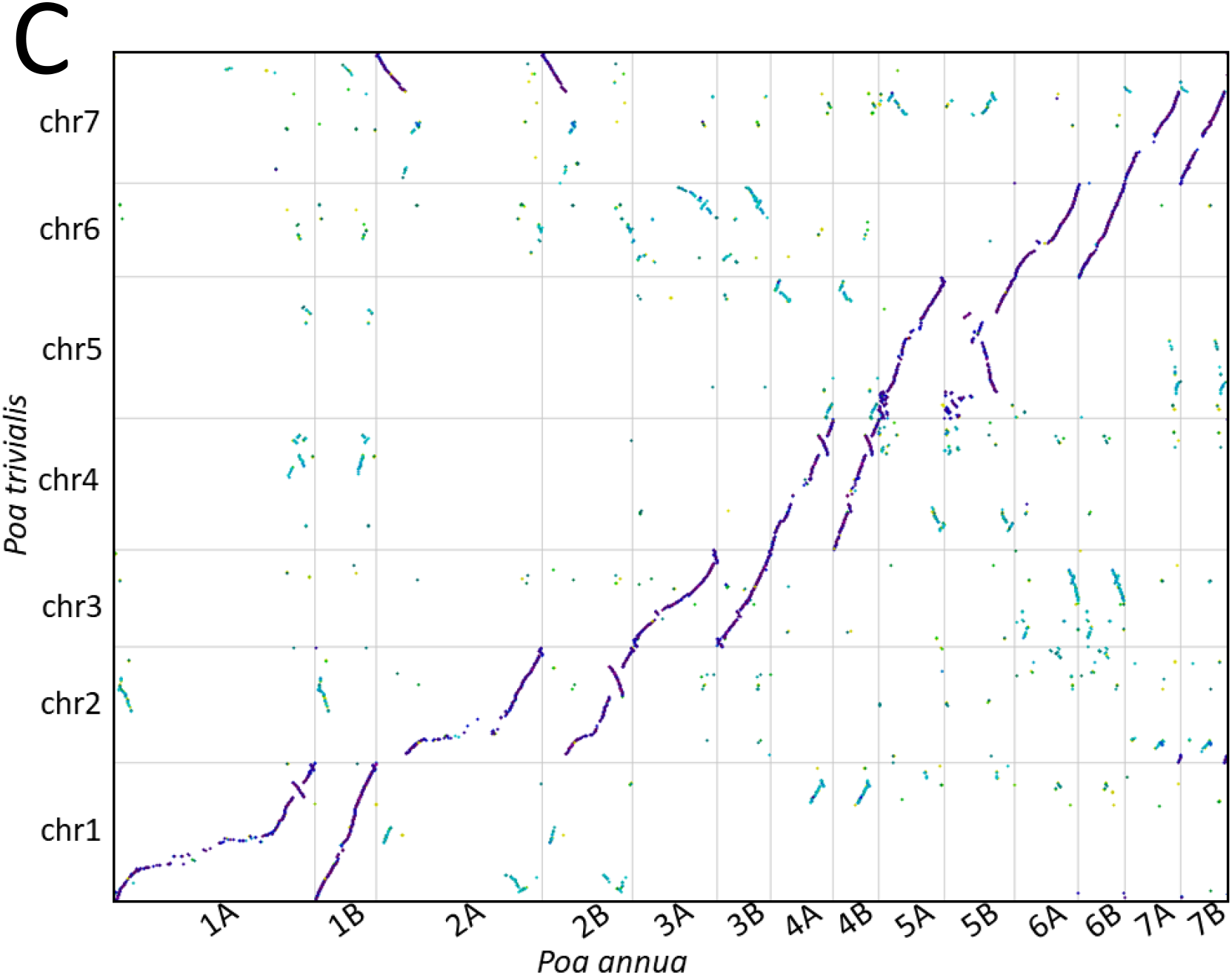
Comparative genomic analysis of *P. trivialis* and other species in the Poaceae family. (A) Synonymous mutation rate (K_s_) analysis between *P. trivialis* and *P. infirma*, *B. dystachyon* and *O. sativa*. (B) Phylogenetic analysis with single copy orthogroups. (C) Syntenic analysis between *P. trivialis* and *P. annua*.

### Population genomics of weedy P. trivialis populations

We surveyed 12 *P. trivialis* populations (Table 3, Figure 3) from agronomic crops throughout the Willamette Valley, a region located in western Oregon where most of the cool-season grass seed crop is grown in the United States. We sampled fields after communication with farmers and crop consultants to identify fields with a known history of infestation with *P. trivialis*, that they considered difficult to manage using common weed control techniques. To genotype populations, we performed reduced-representation sequencing of 10-20 individuals per populations using the NextRAD approach (Russello et al., 2015). We also included an outgroup species (population W2, *P. annua*), 13 herbarium vouchers from the Oregon State University Herbarium that were sampled in the region between 1902 and 2006 (HE), and a *P. trivialis* crop variety that is widely grown in the USA (CR). As expected, the outgroup *P. annua* clearly differentiated from the other groups and separated by PC1 (data not shown), while the crop variety, herbarium vouchers, and weed populations of *P. trivialis* were all closely related and clustered together (Figure 4A). It was possible to visualize that all samples had continuous variation in relatedness, with no clear cluster distinction (Figure 4B). The crop and herbarium samples exhibited rather limited genetic variability. With the same dataset, we performed a population structure analysis using the ADMIXTURE program (Alexander et al., 2009) to further dissect the relatedness among populations. A clear distinction between the outgroup population (*P. annua;* W2) and the rest of the populations was observed (data not shown) with *K*=2. With *K*=5, the relatedness of the crop variety population (CR) with the weed populations remained relatively constant, particularly K0, P2, and individuals in RB1, RB5, and W3. Results from this analysis also suggest that populations are composed of individuals from distinct ancestries, as well as the presence of individuals with multiple ancestries within a population. When the outgroup population was removed from the analysis (Figure S1), the relationship between the CR (crop) population and other weed populations remained relatively constant, with some exceptions, indicating that some of the weed populations had lower relatedness to CR.

**Figure 3.**
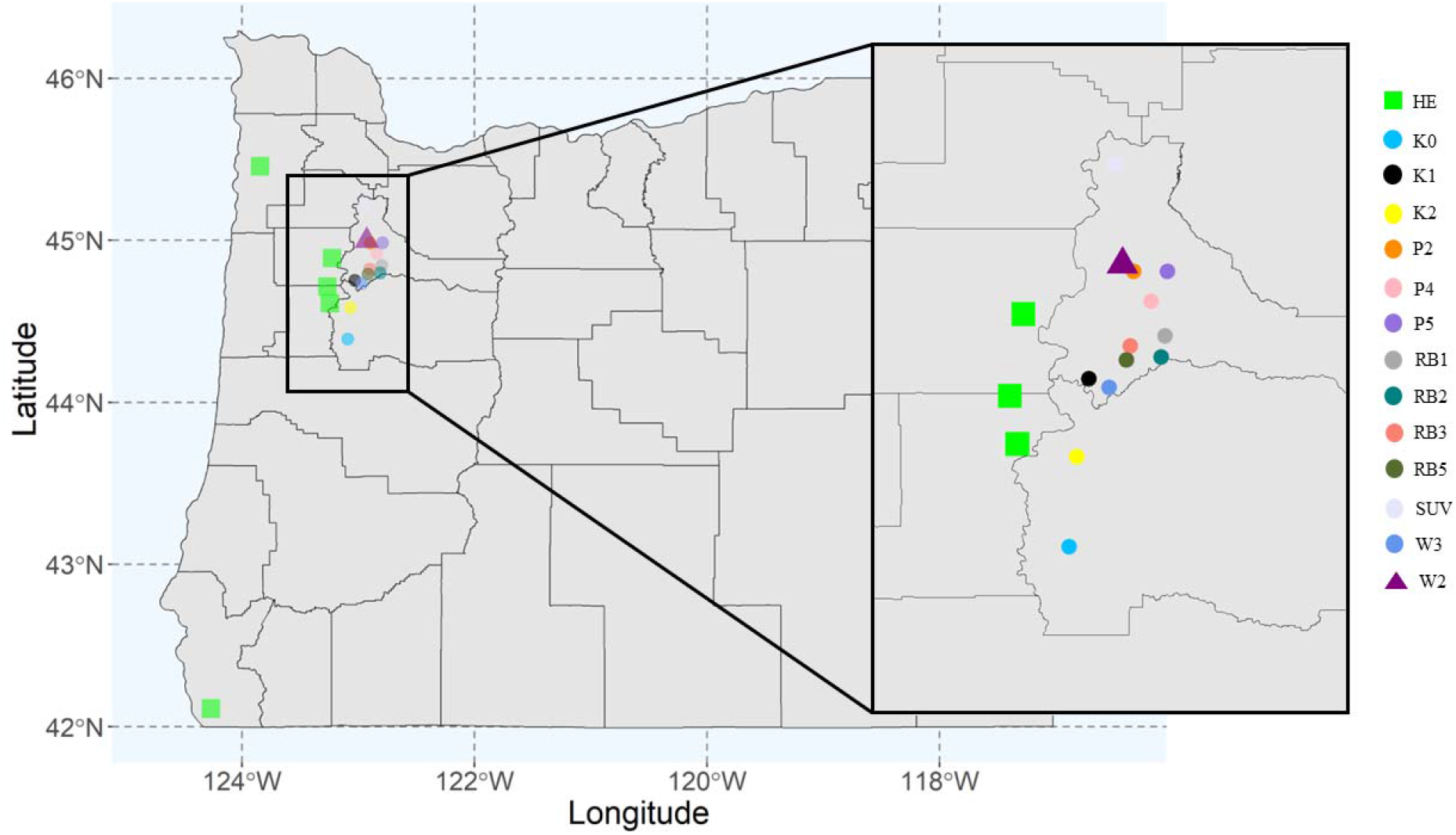
Origin of *Poa trivialis* populations included in this study. Weed populations from agricultural fields (circles, each color corresponds to a different population), herbarium individuals (squares), and an outgroup species *Poa annua* (triangle) were included in the analysis.

**Figure 4.**
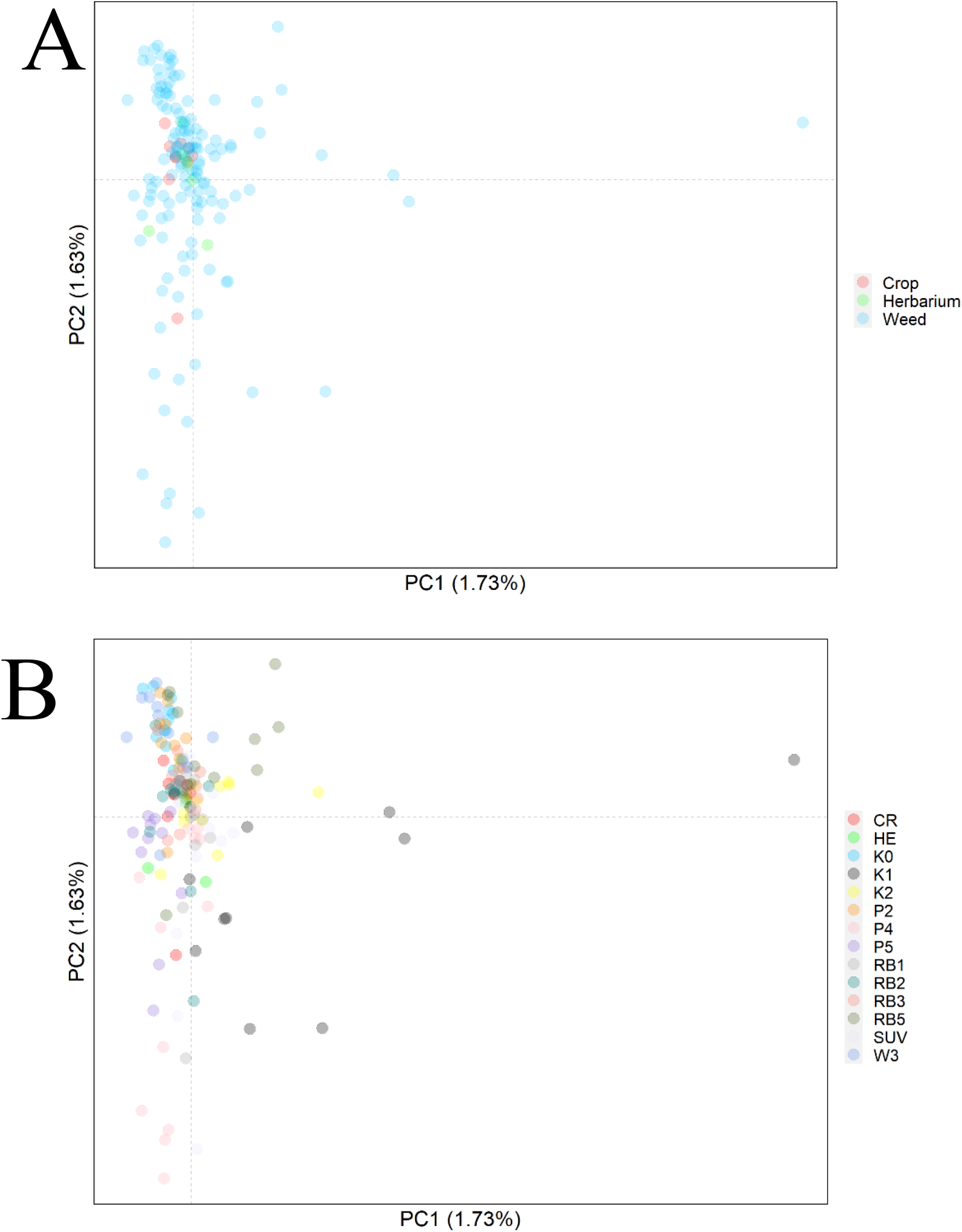
Principal component analysis of *Poa trivialis*. (A) Weed populations, herbarium individuals, and a crop variety were included, resulting in 150 individuals and 73,506 SNPs. (B) Same dataset but colored by population.

**Table 3.** *Poa trivialis* populations included in this study and population genetics estimates.

| Sample<sup>1</sup> | Latitude | Longitude | Sample<br>number | H<sub>E</sub><sup>2</sup> | $\pi$ | F<sub>IS</sub> | Tajima's<br>D |
| --- | --- | --- | --- | --- | --- | --- | --- |
| CR | - | - | **10** | 0.032 | 0.033 | 0.023 | -0.66 |
| HE | - | - | **6** | 0.034 | 0.036 | 0.052 | -0.75 |
| K0 | 44.39 | -123.088 | **11** | 0.030 | 0.031 | 0.009 | -0.55 |
| K1 | 44.752 | -123.028 | **11** | 0.035 | 0.037 | 0.027 | -0.63 |
| K2 | 44.58411 | -123.064 | **11** | 0.032 | 0.034 | 0.024 | -0.92 |
| P2 | 44.98271 | -122.892 | **11** | 0.031 | 0.032 | 0.019 | -0.80 |
| P4 | 44.91874 | -122.839 | **12** | 0.035 | 0.037 | 0.030 | -0.75 |
| P5 | 44.983 | -122.789 | **11** | 0.031 | 0.032 | 0.015 | -0.67 |
| RB1 | 44.84413 | -122.796 | **11** | 0.032 | 0.034 | 0.020 | -0.92 |
| RB2 | 44.7987 | -122.809 | **12** | 0.033 | 0.034 | 0.023 | -0.94 |
| RB3 | 44.8223 | -122.902 | **11** | 0.033 | 0.035 | 0.026 | -0.94 |
| RB5 | 44.79196 | -122.913 | **11** | 0.035 | 0.037 | 0.026 | -0.84 |
| SUV | 45.2137 | -122.951 | **12** | 0.035 | 0.036 | 0.025 | -0.96 |
| W3 | 44.73264 | -122.967 | **11** | 0.032 | 0.034 | 0.011 | -0.56 |
| W2 | 44.99815 | -122.926 | **20** | 0.043 | 0.044 | -0.04 | 0.64 |

<sup>1</sup>CR is a cultivated variety of *Poa trivialis*. HE are herbarium populations from various locations in Oregon. Herbarium vouchers collected in 1919, 1927, 1984, 1996, 1997, and 2001. All other populations are weeds from agricultural fields. <sup>2</sup>H<sub>E</sub>: Expected heterozygosity over polymorphic sites; $\pi$ : nucleotide diversity; F<sub>IS</sub>: Inbreeding coefficient.

A Neighbor-Joining tree was created to further dissect the relationships among *P. trivialis* populations. Results indicated that the herbarium plants clustered together (Figure S2), closer to the crop plants. Three other distinct clusters were observed, each containing a different subset of weedy populations. To understand whether this clustering had geographical patterns, we performed a Mantel test to assess isolation-by-distance. This analysis tests whether genetic differentiation among populations could be explained by their geographical distance from each other, and is calculated by correlating genetic distance and place of origin. We found no significant evidence of isolation-by-distance, indicating that there may be substantial gene flow among populations (Mantel test, *P* = 0.16; Figure S3).

The level of expected heterozygosity (H_E_), nucleotide diversity (π), and inbreeding coefficient (F_IS_) was calculated for each populations using 73,506 SNP’s with the Stacks pipeline (Rivera-Colón & Catchen, 2021), and Tajima’s D with *vcftools* (Danacek et al., 2011). All *P. trivialis* populations exhibited similar H_E_, that ranged from 0.030 to 0.035, while the outgroup (*P. annua*) had a greater H_E_. The π estimate had similar trends, varying from 0.031 to 0.037. Much greater variation was observed for F_IS_, where field-collected populations varied from 0.009 to 0.030, the outgroup had −0.04, and herbarium individuals the largest value (F_IS_ = 0.052). Tajima’s D statistics was negative for all sampled populations, with exception for the outgroup population, indicating an excess of low-frequency alleles. It should be noted that the sequencing data for the outgroup population was aligned to the *P. trivialis* genome, which could have influenced these results because of mapping error.

## Discussion

In this work, we assembled and analyzed the genome of *P. trivialis*, and studied field populations infesting agronomic crops in Oregon. Our assembly of a haploid genome resulted in seven chromosome-scale scaffolds with a high degree of completeness. Further analysis of the *P. trivialis* genome revealed a close genetic relationship with other grass species important for agriculture, such as *P. annua*, *H. vulgare*, and *T. aestivum* and provided insight into genomic modifications that occurred during its evolution. Not surprisingly, the *Poa* genus clustered more with other C_3_ species, compared to the more distantly related C_4_ (i.e., *Setaria viridis*, *Echinochloa crus-galli*, and *Zea mays*). Furthermore, analysis of *P. trivialis* populations revealed limited genetic diversity (π = 0.031-0.037, H_E_ = 0.030-0.035) not only among field-collected individuals, but also compared to herbarium vouchers, and a commonly used crop variety. These values, compared to other outcrossing grass species, such as *Dactylis glomerata* (H_E_ = 0.44-0.59; 50 populations analyzed; >30 individuals per population; Last et al., 2013), *Lolium perenne* (H_E_ = 0.315–0.333; 120 populations analyzed; >50 individuals per population; Balfourier et al., 1998), and *L. rigidum* (0.394–0.424; 50 populations analyzed; >50 individuals per population; Balfourier et al., 1998), further support the limited genetic diversity found in other analyses. Similar results were observed when weedy and cultivated *L. multiflorum* populations were compared, in that both types exhibited similar levels of genetic diversity (Brunharo & Streisfeld, 2022).

The size of the *P. trivialis* genome is in accordance with *k*-mer based analysis and flow cytometry, with a final haploid assembly of 1.35 Gb. As observed in other grass species with large genomes, such as *Z. mays* (Jiao et al., 2017), *P. infirma* (Benson et al., 2023), and *H. vulgare* (Mascher et al., 2017), repetitive elements are abundant in *P. trivialis*, with more than 77% of DNA content represented by various families of transposable elements. Long terminal repeat (LTR) retrotransposons from the *Copia* and *Gypsy* superfamilies were dominant in *P. trivialis* (>46%), as typically observed in other plant species, with content as large as 70% (Schnable et al., 2009). Over 58k gene models were identified using protein-homology and isoform-sequencing (IsoSeq), of which 65% were functionally annotated using public databases. This low number of functionally annotated genes can be explained by the divergence of *P. trivialis* from other species available in the Uniprot database with our chosen similarity cutoffs.

A total of 12 *P. trivialis* field populations were included in the study. The specific agronomic region where *P. trivialis* is typically grown for seed is limited to central Oregon, in Jefferson County, with approximately 450 ha total in the state, and none of the populations we sampled originated from that region (except for the crop variety). The Willamette Valley of Oregon is the most important cool-season grass seed region in the United States, with typical crops perennial ryegrasss, annual ryegrass, and tall fescue, often in rotation with *Trifolium spp*. Most of the populations included in this study were sampled from those agronomic crop fields.

Our PCA indicated that the *P. trivialis* field populations had no distinct patterns of genetic structure compared to the crop variety (CR; Figure 4A and 4B), suggesting a close genetic relationship. The lack of population structure between crop vs. weedy genotypes in *P. trivialis* observed in the PCA could be explained by three reasons. First, the weed populations could have originated from CR or a similar crop variety, and later spread throughout the region, explaining the close genetic relationship with CR. The founding variety could have been an adapted variety for the region and spread with contaminated seed lots. The greater genetic variation observed in the weedy field populations in Figure 4A could be due to an ongoing process of de-domestication or gene flow. Second, given the populations were collected from agricultural fields, it is possible that human-facilitated, recurrent gene flow is ongoing in the region. In agricultural fields, it is common to observe long-distance gene flow facilitated by contaminated seed lots and farm equipment, as well as pollen-mediated gene flow in this open-pollinated outcrossing species. Third, it is also possible that the weedy field populations are, in reality, descendent from new, unintentional introductions of *P. trivialis* that dated >100 years ago, as herbarium individuals seem to have close relatedness with some of the crop and weed populations. This hypothesis is further supported by the Neighbor-Joining analysis, where population CR is closely related to the herbaria plants. The oldest herbarium individual included in the analysis was collected in 1919, although in our collection we had individuals from 1902 that were excluded from the analysis due to the large number of missing SNPs.

It is likely that all aspects suggested above, to a certain extent, played a role in the spread of *P. trivialis*, and explain the close relationship between field populations and crop individuals. This is supported by population structure analysis, in that CR and HE individuals exhibited relatedness with some of the weed populations. Overall, this analysis suggests that weedy *P. trivialis* populations have a close relationship with the crop variety. It is unclear, however, whether weed populations descended from the crop varieties and are now experiencing a process of de-domestication, or whether the crop variety originated from a weed population with promising commercial value. Early introductions of *P. trivialis* could be attributed to soil used as ship ballast, as indicated in one of the herbarium vouchers analyzed (voucher number ORE15609, from 1902, of the Oregon State University Herbarium).

The overall low levels of inbreeding, π, and H_E_ observed (Table 3) further supports the assumption of high levels of gene flow among populations. The low level of inbreeding is expected given no clear structuring was observed in the PCA, and similar results were obtained elsewhere for invasive plant species with high levels of gene flow (e.g., *Rosa rugosa*; Kelager et al., 2013). The fact that Tajima’s D values were negative and the low diversity measures for all weedy populations may indicate that populations have an excess of rare alleles, likely because of constant bottlenecks (e.g., local extinction from plowing, herbicide followed by rapid recolonization) imposed by agricultural practices in the region.

The *P. trivialis* genome reported in this research will be an important tool for the scientific community. Because *P. trivialis* has colonized many environments, this species could be used as a model to understand the evolution of traits and the genetic underpinnings of phenotypic plasticity. In fact, other species in the *Poa* genus (Benson et al., 2021; Grime & Mackey, 2002; Molina-Montenegro et al., 2016) also exhibit wide distribution around the world that is likely facilitated by their plasticity. One of the most important components of a weed management plan is to prevent new weed introductions and to limit dispersal. Although seed lot contamination has historically been the prominent process for gene flow, other mechanisms cannot be ruled out, such as farm implement contamination and wind-mediated gene flow. The *P. trivialis* genome could also be used as a tool to identify genetic markers for the quick detection of contaminated seed lots, especially in situations where visual seed inspection and discrimination is difficult due to similarities between crop and weed seed, such as weedy *P. trivialis* seed contamination in commercial seed lots of *P. annua* or *P. pratensis*. Such methods have been used for detecting seed from the noxious *Amaranthus palmeri* weed in seed mixes with other desirable *Amaranthus spp.* (Brusa et al., 2021). Identifying contamination of weedy biotypes of *P. trivialis* in cultivated *P. trivialis* would be challenging, given their relatedness observed in our studies. However, identifying the genetic bases of weedy traits could circumvent this challenge, targeting specific genes or genomic regions for diagnosis of seed lot contamination. Weeds typically show high tolerance for biotic and abiotic stresses, and community efforts have been initiated to generate genomic resources for weeds, with the goal of identifying the genetic architecture of adaptation. Given *P. trivialis* has colonized many environments, dissecting the genetic mechanisms of adaptation could help guide breeding efforts not only for this species, but also for other cereal crops to improve stress tolerance in increasingly fluctuating agricultural ecosystems.

## Material and Methods

### Genome size quantification

Tissue from the youngest fully expanded leaf of a healthy *P. trivialis* individual was sampled, immediately frozen in liquid nitrogen, and submitted for genome size determination. A commercial kit from BD Biosciences (Franklin Lakes, NJ) was used to prepare the leaf tissue, and analysis in a BD FACS Calibur flow cytometer at Lifeasible (www.lifeasible.com). The genome size of *P. trivialis* was determined by comparing to *Solanum lycopersicum* as a known reference at 1C = 1.03 pg. We also quantified the *P. trivialis* genome computationally. Jellyfish (Marçais & Kingsford, 2011) was used to count the number of *k*-mers of length 21 in all HiFi reads, and generate a *k*-mer histogram, followed by plotting and quantification of genome size and heterozygosity with GenomeScope 2.0 (Ranallo-Benavidez et al., 2020).

### Genome Sequencing and Assembly

Leaf tissue for high molecular weight DNA and RNA extraction was collected from a single individual, and genetic material was extracted with a commercial kit (Wizard HMW DNA Extraction and RNeasy Plant Mini Kit). Approximately 68 Gb of high-accuracy (predicted accuracy >99%) PacBio long read data was generated, equivalent to 25 × coverage based on flow cytometry estimates of genome size. A Hi-C Plant Kit (Phase Genomics, Seattle, WA) was used to create a proximity ligation library that included four restriction enzymes (DpnII, DdelI, HinFI, and MseI). The Hi-C library was sequenced using an Illumina Novaseq 6000, producing 114 Gb of data.

We used *hifiasm* 0.16.1-r375 (Cheng et al., 2021) to generate a haploid genome assembly of *P. trivialis*. After parameter optimization based on contiguity of the primary assembly and BUSCO scores (v.5.4.3) (Manni et al., 2021), we implemented *hifiasm* with Hi-C integration using the raw short-read fastq files with default parameters (except *--primary*, *-s 0.1*, and *-l2*). After completion of an initial assembly, Hi-C data was processed following the Arima Genomics pipeline (https://github.com/ArimaGenomics/mapping_pipeline). Briefly, raw forward and reverse short-reads DNA were aligned separately to the draft genome obtained with the previous step with *bwa-mem* 0.7.17-r1188 (Li & Durbin, 2009) and sorted with *Samtools* (Li et al., 2009). Chimeric reads were filtered out with an open source script (https://github.com/ArimaGenomics/mapping_pipeline/blob/master/filter_five_end.pl), and forward and reverse reads were then combined (https://github.com/ArimaGenomics/mapping_pipeline/blob/master/two_read_bam_combiner.pl). PCR duplicates were marked and discarded with *PicardTools* (http://broadinstitute.github.io/picard). With the pre-processed Hi-C data and the draft genome, we utilized YaHS (Zhou et al., 2022) for scaffolding (included the flag *-e GATC,TNA,ANT,TA* to specify the restriction enzymes used in the Hi-C library preparation). We performed a manual curation of the genome with Juicebox (Durand et al., 2016) to fix mis-assemblies. Finally, we performed a whole-genome alignment to *P. annua* with *minimap2* (v.2.23-r1111) (Li, 2018) to assign chromosome numbers to *P. trivialis*.

### Repeat and Gene Annotation

The gene space of *P. trivialis* was annotated with two IsoSeq libraries using RNA extracted from leaf and root tissues, that was multiplexed and sequenced with a single PacBio SMRT cell 8M at the Center for Quantitative Life Science at Oregon State University. The data was processed with the IsoSeq v3 pipeline (https://github.com/PacificBiosciences/IsoSeq). Briefly, full-length transcripts were demultiplexed and had the sequencing primers removed with the *lima* algorithm, following by the poly-A trimming and concatemer identification and removal with the *refine* program. Finally, the *cluster* algorithm was used to cluster the isoforms. *Minimap2* (Li, 2018) was used to align the clustered full-length transcripts to the reference genome, following by collapsing of isoforms from leaf and root tissue with *TAMA* (*-d* merge_dup *-x* no_cap *-a* 100 *-z* 100 *-sj* no_priority *-sjt* 10) (Kuo et al., 2020).

Transposable elements (TE) were annotated with *Extensive De-novo TE Annotator* (EDTA) (Ou et al., 2019) using our reference genome and the transcriptome generated in the previous step. This step included the use of *RepeatModeler* (Flynn et al., 2020) to identify transposable elements missed by the EDTA algorithm, generating a non-redundant TE library.

The final step in the annotation pipeline was to perform three rounds of *Maker* with default parameters (Campbell et al., 2014) to identify the genomic features of *P. trivialis*. We used the protein sequences of *Brachypodium distachyon*, *Triticum aestivum*, and *Hordeum vulgare* from EnsemblPlants (v. 107.4) (Cunningham et al., 2021), as well as the TE library and transcriptome generated in previous steps. The gene models generated in the first run of *Maker* were used for training gene prediction using *SNAP* (Korf, 2004) and *Augustus* (Stanke et al., 2006).

### Genome statistics and comparative genomics

We assessed the genome completeness with BUSCO (v5.4.3) (Manni et al., 2021) with the *Poales* database. Assembly statistics were computed with the *seqstat* program from *GenomeTools* (http://genometools.org/). Quast v5.2.0 (Gurevich et al., 2013) was used to quantify additional assembly statistics such as L_50_, and GC content. The phylogenetic relationship of *P. trivialis* was assessed in comparison to *P. supina*, *P. infirma*, *Zea mays*, *T. aestivum*, *H. vulgare*, *B. distachyon*, *Oryza sativa*, *Echinochloa crus-galli*, and *Setaria viridis*. We used *OrthoFinder* v2.5.4 (Emms & Kelly, 2019) to identify single-copy orthogroups to construct, including the *-msa* option for multiple sequence alignment with a built-in MAFFT (Katoh et al., 2002). A phylogenetic analysis was conducted with the algorithm *RAxML-NG* (Kozlov et al., 2019). Phylogenetic tree plotting was performed with custom scripts in R, using the packages *ggtree* (Xu et al.) and *treeio* (Wang et al., 2019).

Syntenic dotplots were generated in CoGe (https://genomevolution.org/coge/) to obtain an overview of the structural variation between *P. trivialis* and *P. annua* using the SynMap tool. In addition, the synonymous mutation rate (K_s_) was calculated for protein coding pairs to estimate divergence time, based on a mutation rate of 5.76174 x 10^-9^ (De La Torre et al., 2017), compared to *P. infirma*, *B. distachyon*, and *O. sativa*.

### Population genomics analysis

We sampled 12 populations of *P. trivialis* from agricultural fields in the Willamette Valley region of Oregon (Figure 3). The fields were identified with the assistance of crop consultants and extension personnel, who identified the populations that were difficult to manage utilizing conventional weed control techniques, particularly herbicides.

At each field, leaf tissue from 15-20 individuals were collected when plants were at the flowering stage, and kept in dry ice until long-term storage in a −80 C freezer upon return to the laboratory. Seeds were also collected for greenhouse studies. We also obtained the cultivated variety “Quasar “ of *P. trivialis* that is commonly grown for seed production, as well as a *P. annua* population used as an outgroup. Each collected individual was genotyped with the nextRAD (Nextera-tagmented, Reductively-Amplified DNA), as follows. DNA was extracted using a commercial kit (Wizard Genomic DNA Purification Kit, Promega, Madison, WI). Libraries were produced by first fragmenting 75 ng DNA with the Nextera reagent (Illumina, Inc), and ligation of short adapter sequences to fragments. Fragmented DNA was amplified for 27 cycles at 74 degrees, with one of the primers matching the adapter and extending 10 nucleotides into the genomic DNA with a selective sequence (Russello et al., 2015). The nextRAD libraries were sequenced on a Novaseq 6000 with the S1 flow cell and 122 cycles.

Single-ended reads were pre-processed with the *HTStream* suite (https://github.com/s4hts/HTStream), and included contaminant screening, adapter and “N” trimming, removal of bases with quality below Q20, and keeping reads longer than 50 bp. Reads passing quality control were aligned to the *P. trivialis* reference genome with *bwa-mem* (Li & Durbin, 2009) and sorted with *Samtools* (Li et al., 2009). The *ref_map.pl* wrapper from Stacks v2.52 (Rochette et al., 2019) was used to process alignment files and identify single nucleotide polymorphisms (SNPs), and the program *populations* to generate population-level statistics. For all population comparisons, we maintained a minimum of 50% of samples per population *(--min-samples-per-pop 0.5*), and minor-allele frequency of 5% (*--min-maf 0.05*), and only identified a single SNP per locus (*--write-single-snp*). When we excluded the outgroup population from the analysis, we increase the missing threshold to 80%. The *vcf* files produced by the populations program were used in all downstream analysis unless otherwise noted. The population genetics estimates expected heterozygosity (H_E_), nucleotide diversity (π), and inbreeding coefficient (F_IS_) were obtained with the *populations* module of Stacks. For each population, the Tajima’s D statistics was computed with *vcftools* in 10 kb sliding windows.

We performed a principal component analysis (PCA) to obtain an overall picture of the genetic background of field populations. This is a model-free method that helps to visually represent and understand genetic structure (McVean, 2009). Two independent analyses were performed with and without an outgroup. The analysis with an outgroup population included 11,776 SNPs and 179 individuals that passed the filtering criteria, while the analysis without the outgroup had 73,506 SNPs and 152 individuals. To complement the PCA, we further analyzed the population structure of *P. trivialis* populations with ADMIXTURE v0.1.13 (Alexander et al., 2009), and included an outgroup population, with *K* varying from 2-5. The Neighbor-Joining tree was performed by obtaining pairwise genetic distances with the R package *adegent* (Jombart, 2008), estimation with the *nj* function from the *ape* package (Paradis et al., 2004), and plotting with *ggtree*. Isolation-by-distance analysis was performed with the *dist.genpop* function (to obtain genetic distances) from the *adegenet* R package, and the Geographic Distance Matrix Generator tool for obtaining geographic distances (Ersts, 2023), and assessed with the *mantel.randtest* function with 10,000 permutations.

## Acknowledgements

We would like to thank Dr. Aaron Liston for his assistance to identify and sample *P. trivialis* tissue from accessions available at the Oregon State University Herbarium. Funding for this project was provided by the College of Agricultural Sciences at The Pennsylvania State University and Oregon State University.

## Data Statement

The reference genome and related files (i.e., protein, transcript, GFF) are available at the International Weed Genomics Consortium (https://www.weedgenomics.org/). Raw data for population genomics studies are available at Sequence Read Archive BioProject PRJNA1034650.

---

**Figure S1.**
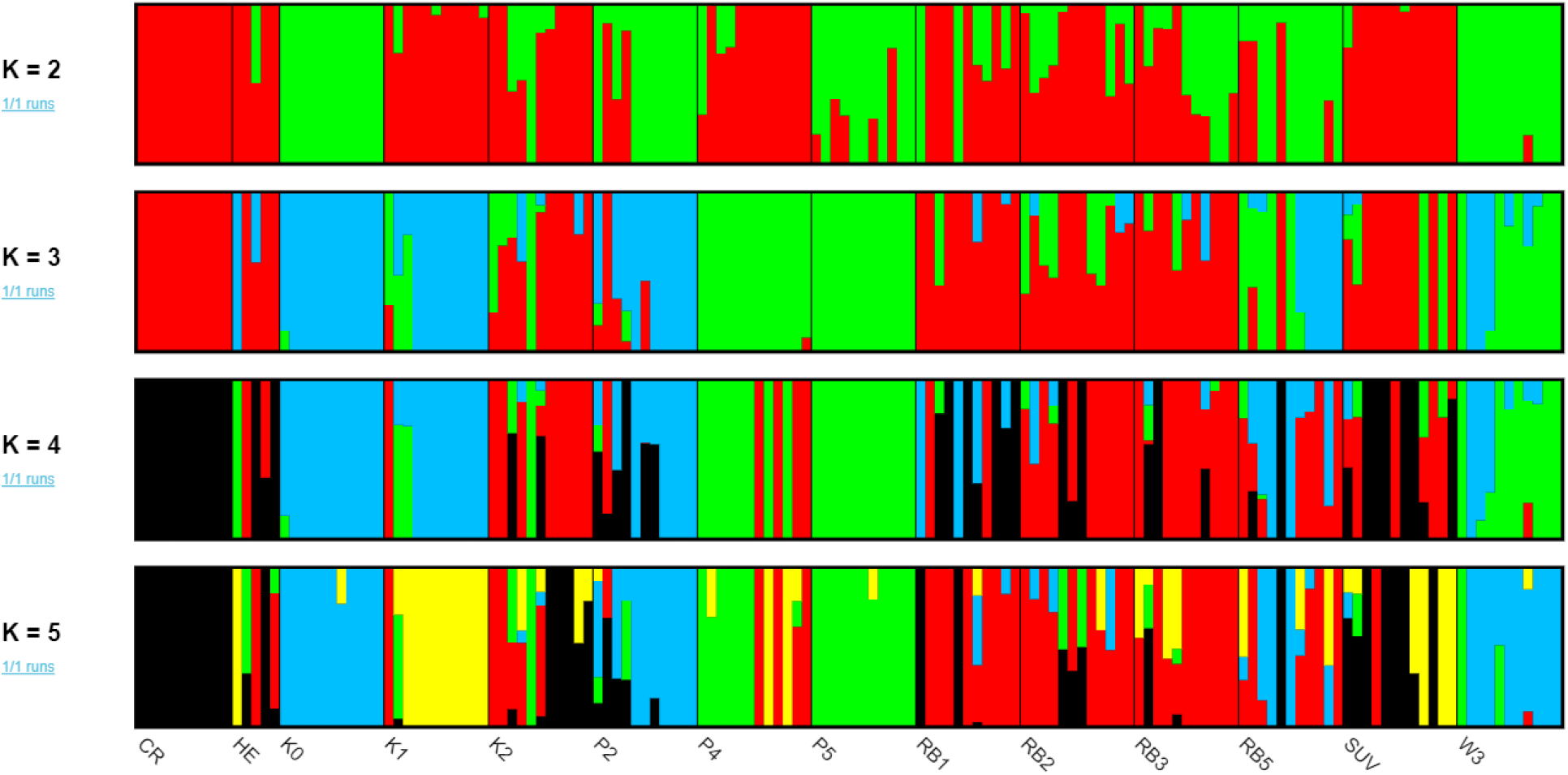
ADMIXTURE analysis estimates individual ancestries and gene flow between herbarium vouchers (HE), field populations, and a cultivated variety (CR) of *P. trivialis*.

**Figure S2.**
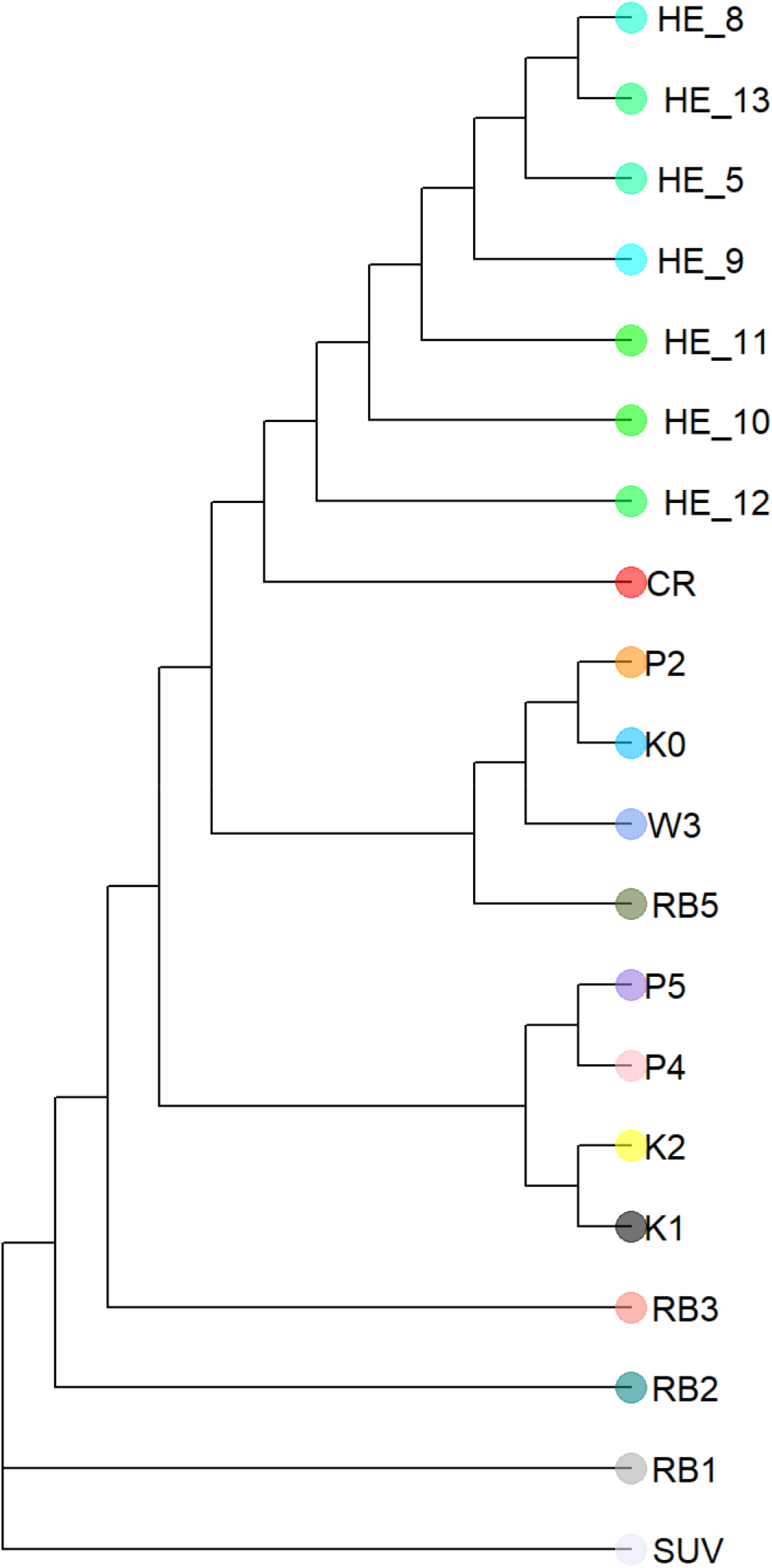
Rooted Neighbour joining tree of *P. trivialis* populations.

**Figure S3.**
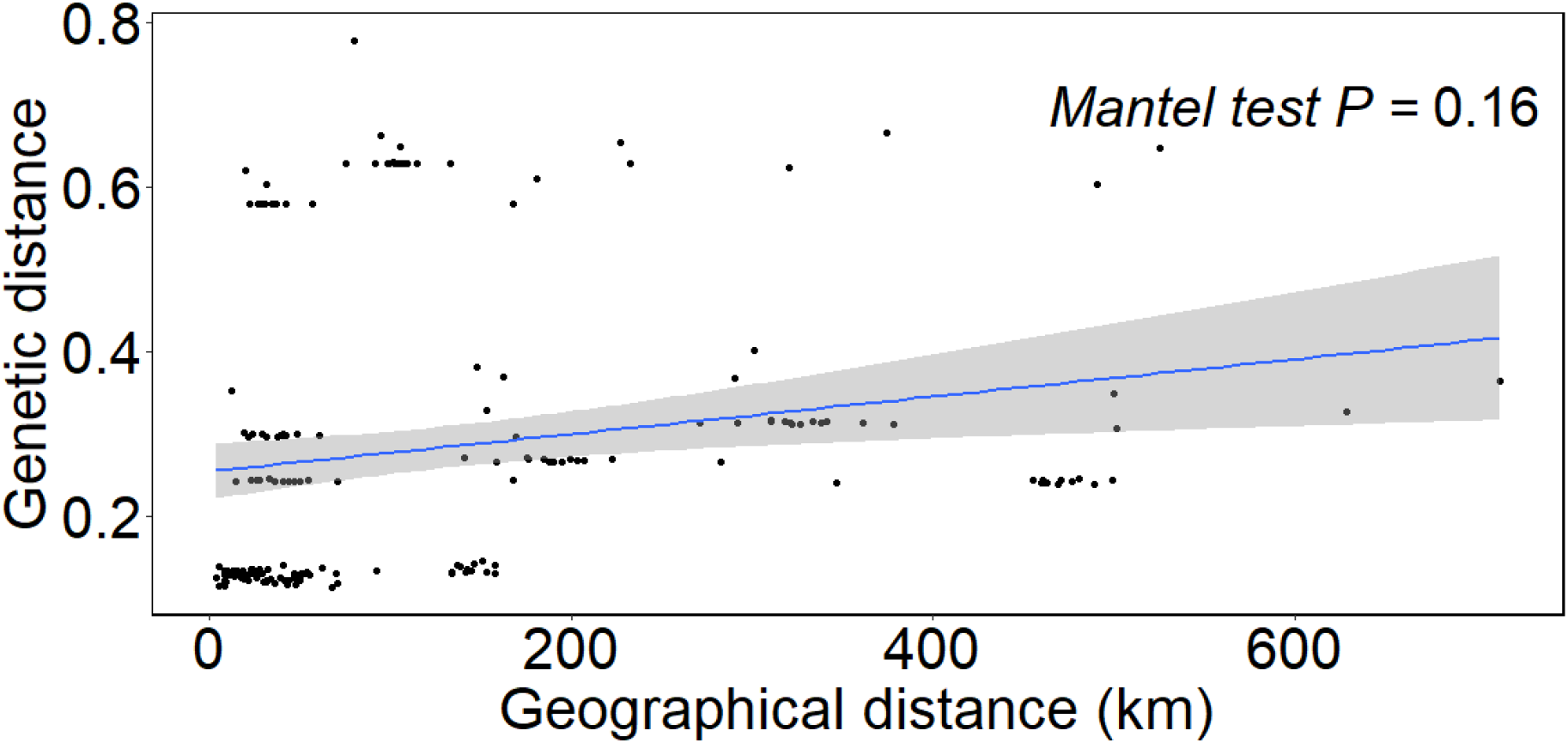
Isolation-by-distance plot with pairwise genetic and geographic distance between *P. trivialis* individuals. Mantel test was performed with 10,000 simulations (*P*=0.16)

## References

Ahmed, M. K., Jelenkovic, G., Dickson, W. R., & Funk, C. R. (1972). CHROMOSOME MORPHOLOGY OF POA TRIVIALIS L. Canadian Journal of Genetics and Cytology, 14(2), 287–291. 10.1139/g72-036

Alexander, D. H., Novembre, J., & Lange, K. (2009). Fast model-based estimation of ancestry in unrelated individuals. Genome Research, 19(9), 1655–1664. 10.1101/gr.094052.109

Balfourier, F., Charmet, G., Ravel, C. (1998) Genetic differentiation within and between natural populations of perennial and annual ryegrass (Lolium perenne and L. rigidum). Heredity 81: 100–110.

Benson, C. W., Mao, Q., & Huff, D. R. (2021). Global DNA methylation predicts epigenetic reprogramming and transgenerational plasticity in Poa annua L. Crop Science, 61(5), 3011–3022. 10.1002/csc2.20337

Benson, C. W., Sheltra, M. R., Maughan, P. J., Jellen, E. N., Robbins, M. D., Bushman, B. S., Patterson, E. L., Hall, N. D., & Huff, D. R. (2023). Homoeologous evolution of the allotetraploid genome of Poa annua L. BMC Genomics, 24(1), 350. 10.1186/s12864-023-09456-5

Brunharo, C. A. C. G., Streisfeld, M. A. (2022) Multiple evolutionary origins of glyphosate resistance in Lolium multiflorum. Evolutionary Applications, 15, 316–329. 10.1111/eva.13344

Brusa, A., Patterson, E. L., Gaines, T. A., Dorn, K., Westra, P., Sparks, C. D., & Wyse, D. (2021). A needle in a seedstack: an improved method for detection of rare alleles in bulk seed testing through KASP. Pest Management Science, 77(5), 2477–2484. 10.1002/ps.6278

Campbell, M. S., Holt, C., Moore, B., & Yandell, M. (2014). Genome Annotation and Curation Using MAKER and MAKER-P. Current Protocols in Bioinformatics, 48(1), 4.11.11–14.11.39. 10.1002/0471250953.bi0411s48

Cheng, H., Concepcion, G. T., Feng, X., Zhang, H., & Li, H. (2021). Haplotype-resolved de novo assembly using phased assembly graphs with hifiasm. Nature Methods, 18(2), 170–175. 10.1038/s41592-020-01056-5

Cunningham, F., Allen, J. E., Allen, J., Alvarez-Jarreta, J., Amode, M R., Armean, Irina M., Austine- Orimoloye, O., Azov, Andrey G., Barnes, I., Bennett, R., Berry, A., Bhai, J., Bignell, A., Billis, K., Boddu, S., Brooks, L., Charkhchi, M., Cummins, C., Da Rin Fioretto, L., … Flicek, P. (2021). Ensembl 2022. Nucleic Acids Research, 50(D1), D988–D995. 10.1093/nar/gkab1049

Danacek, P., Auton, A., Abecasis, G., Albers, C., Banks, E., DePristo M., Handsaker, R., Lunter, G., Marth, G., Sherry, S., McVean, G., Durbin, R., 1000 Genomes Project Analysis Group (2011) The variant call format and VCFtools. Bioinformatics, 27(15), 2156–2158. 10.1093/bioinformatics/btr330

De La Torre, A. R., Li, Z., Van de Peer, Y., & Ingvarsson, P. K. (2017). Contrasting Rates of Molecular Evolution and Patterns of Selection among Gymnosperms and Flowering Plants. Molecular Biology and Evolution, 34(6), 1363–1377. 10.1093/molbev/msx069

Durand, N. C., Robinson, J. T., Shamim, M. S., Machol, I., Mesirov, J. P., Lander, E. S., & Aiden, E. L. (2016). Juicebox Provides a Visualization System for Hi-C Contact Maps with Unlimited Zoom. Cell Systems, 3(1), 99–101. 10.1016/j.cels.2015.07.012

Emms, D. M., & Kelly, S. (2019). OrthoFinder: phylogenetic orthology inference for comparative genomics. Genome Biology, 20(1), 238. 10.1186/s13059-019-1832-y

Ersts, P. J. (2023). Geographic Distance Matrix Generator(version 1.2.3). . American Museum of Natural History, Center for Biodiversity and Conservation. . http://biodiversityinformatics.amnh.org/open_source/gdmg

Flynn, J. M., Hubley, R., Goubert, C., Rosen, J., Clark, A. G., Feschotte, C., & Smit, A. F. (2020). RepeatModeler2 for automated genomic discovery of transposable element families. Proceedings of the National Academy of Sciences, 117(17), 9451–9457. doi:10.1073/pnas.1921046117

George, W. M.-W. (1990). Control of Roughstalk Bluegrass (Poa trivialis) with Fenoxaprop in Perennial Ryegrass (Lolium perenne) Grown for Seed. Weed Technology, 4(2), 250–257. http://www.jstor.org/stable/3987069

Grime, J. P., & Mackey, J. M. L. (2002). The role of plasticity in resource capture by plants. Evolutionary Ecology, 16(3), 299–307. 10.1023/A:1019640813676

Gurevich, A., Saveliev, V., Vyahhi, N., & Tesler, G. (2013). QUAST: quality assessment tool for genome assemblies. Bioinformatics, 29(8), 1072–1075. 10.1093/bioinformatics/btt086

Haggar, R. J. (1971). The significance and control of Poa trivialis in ryegrass pastures. Grass and Forage Science, 26(3), 117–122. 10.1111/j.1365-2494.1971.tb00652.x

Haggar, R. J. (1979). Competition between Lolium perenne and Poa trivialis during establishment. Grass and Forage Science, 34(1), 27–36. 10.1111/j.1365-2494.1979.tb01444.x

Haug-Baltzell, A., Stephens, S. A., Davey, S., Scheidegger, C. E., & Lyons, E. (2017). SynMap2 and SynMap3D: web-based whole-genome synteny browsers. Bioinformatics, 33(14), 2197–2198. 10.1093/bioinformatics/btx144

Heap, I. (2022). International Herbicide-Resistant Weed Database. Retrieved 08/10/2022 from https://weedscience.org/Home.aspx

Jiao, Y., Peluso, P., Shi, J., Liang, T., Stitzer, M. C., Wang, B., Campbell, M. S., Stein, J. C., Wei, X., Chin, C.-S., Guill, K., Regulski, M., Kumari, S., Olson, A., Gent, J., Schneider, K. L., Wolfgruber, T. K., May, M. R., Springer, N. M., Ware, D. (2017). Improved maize reference genome with single-molecule technologies. Nature, 546(7659), 524–527. 10.1038/nature22971

Katoh, K., Misawa, K., Kuma, K. i., & Miyata, T. (2002). MAFFT: a novel method for rapid multiple sequence alignment based on fast Fourier transform. Nucleic Acids Research, 30(14), 3059–3066. 10.1093/nar/gkf436

Kelager, A., Pedersen, J. S., & Bruun, H. H. (2013). Multiple introductions and no loss of genetic diversity: invasion history of Japanese Rose, Rosa rugosa, in Europe. Biological Invasions, 15(5), 1125–1141. 10.1007/s10530-012-0356-0

Korf, I. (2004). Gene finding in novel genomes. BMC Bioinformatics, 5(1), 59. 10.1186/1471-2105-5-59

Kozlov, A. M., Darriba, D., Flouri, T., Morel, B., & Stamatakis, A. (2019). RAxML-NG: a fast, scalable and user-friendly tool for maximum likelihood phylogenetic inference. Bioinformatics, 35(21), 4453–4455. 10.1093/bioinformatics/btz305

Kuo, R. I., Cheng, Y., Zhang, R., Brown, J. W. S., Smith, J., Archibald, A. L., & Burt, D. W. (2020). Illuminating the dark side of the human transcriptome with long read transcript sequencing. BMC Genomics, 21(1), 751. 10.1186/s12864-020-07123-7

Jombart, T. (2008). adegenet: a R package for the multivariate analysis of genetic markers. Bioinformatics, 24(11), 1403–1405. 10.1093/bioinformatics/btn129

Levy, M. (1998). Poa trivialis contamination: An increase in testing standards would benefit superintendents. U. USGA Green Sec. Rec., 36, 13–14.

Last, L., Widmer, F., Fjellstad, W., Stoyanova, S., Kolliker, R. (2013) Genetic diversity of natural orchardgrass (Dactylis glomerata L.) populations in three regions in Europe. BMC Genetics 14:e102.

Li, H. (2018). Minimap2: pairwise alignment for nucleotide sequences. Bioinformatics, 34(18), 3094–3100. 10.1093/bioinformatics/bty191

Li, H., & Durbin, R. (2009). Fast and accurate short read alignment with Burrows–Wheeler transform. Bioinformatics, 25(14), 1754–1760. 10.1093/bioinformatics/btp324

Li, H., Handsaker, B., Wysoker, A., Fennell, T., Ruan, J., Homer, N., Marth, G., Abecasis, G., Durbin, R., & Subgroup, G. P. D. P. (2009). The Sequence Alignment/Map format and SAMtools. Bioinformatics, 25(16), 2078–2079. 10.1093/bioinformatics/btp352

Manni, M., Berkeley, M. R., Seppey, M., Simão, F. A., & Zdobnov, E. M. (2021). BUSCO Update: Novel and Streamlined Workflows along with Broader and Deeper Phylogenetic Coverage for Scoring of Eukaryotic, Prokaryotic, and Viral Genomes. Molecular Biology and Evolution, 38(10), 4647–4654. 10.1093/molbev/msab199

Mao, Q., & Huff, D. R. (2012). The Evolutionary Origin of Poa annua L. Crop Science, 52(4), 1910–1922. 10.2135/cropsci2012.01.0016

Marçais, G., & Kingsford, C. (2011). A fast, lock-free approach for efficient parallel counting of occurrences of k-mers. Bioinformatics, 27(6), 764–770. 10.1093/bioinformatics/btr011

Mascher, M., Gundlach, H., Himmelbach, A., Beier, S., Twardziok, S. O., Wicker, T., Radchuk, V., Dockter, C., Hedley, P. E., Russell, J., Bayer, M., Ramsay, L., Liu, H., Haberer, G., Zhang, X.-Q., Zhang, Q., Barrero, R. A., Li, L., Taudien, S., … Stein, N. (2017). A chromosome conformation capture ordered sequence of the barley genome. Nature, 544(7651), 427–433. 10.1038/nature22043

McVean, G. (2009). A Genealogical Interpretation of Principal Components Analysis. PLOS Genetics, 5(10), e1000686. 10.1371/journal.pgen.1000686

Molina-Montenegro, M. A., Galleguillos, C., Oses, R., Acuña-Rodríguez, I. S., Lavín, P., Gallardo-Cerda, J., Torres-Díaz, C., Diez, B., Pizarro, G. E., & Atala, C. (2016). Adaptive phenotypic plasticity and competitive ability deployed under a climate change scenario may promote the invasion of Poa annua in Antarctica. Biological Invasions, 18(3), 603–618. 10.1007/s10530-015-1033-x

Ou, S., Su, W., Liao, Y., Chougule, K., Agda, J. R. A., Hellinga, A. J., Lugo, C. S. B., Elliott, T. A., Ware, D., Peterson, T., Jiang, N., Hirsch, C. N., & Hufford, M. B. (2019). Benchmarking transposable element annotation methods for creation of a streamlined, comprehensive pipeline. Genome Biology, 20(1), 275. 10.1186/s13059-019-1905-y

Paradis, E., Claude, J., & Strimmer, K. (2004). APE: Analyses of Phylogenetics and Evolution in R language. Bioinformatics, 20(2), 289–290. 10.1093/bioinformatics/btg412

Phillips, A. R., Seetharam, A. S., AuBuchon-Elder, T., Buckler, E. S., Gillespie, L. J., Hufford, M. B., Llaca, V., Romay, M. C., Soreng, R. J., Kellogg, E. A., & Ross-Ibarra, J. (2022). A happy accident: a novel turfgrass reference genome. bioRxiv, 2022.2003.2008.483531. 10.1101/2022.03.08.483531

Rajasekar, S., Fei, S.-h., & Christians, N. E. (2006). Analysis of Genetic Diversity in Rough Bluegrass Determined by RAPD Markers. Crop Science, 46(1), 162–167. 10.2135/cropsci2005.04-0008

Ranallo-Benavidez, T. R., Jaron, K. S., & Schatz, M. C. (2020). GenomeScope 2.0 and Smudgeplot for reference-free profiling of polyploid genomes. Nature Communications, 11(1), 1432. 10.1038/s41467-020-14998-3

Rivera-Colón, A. G., & Catchen, J. (2021). Population genomics analysis with RAD, reprised: Stacks 2. bioRxiv, 2021.2011.2002.466953. 10.1101/2021.11.02.466953

Rochette, N. C., Rivera-Colón, A. G., & Catchen, J. M. (2019). Stacks 2: Analytical methods for paired-end sequencing improve RADseq-based population genomics. Molecular Ecology, 28(21), 4737–4754. 10.1111/mec.15253

Russello, M. A., Waterhouse, M. D., Etter, P. D., & Johnson, E. A. (2015). From promise to practice: pairing non-invasive sampling with genomics in conservation. PeerJ, 3, e1106. 10.7717/peerj.1106

Schnable, P. S., Ware, D., Fulton, R. S., Stein, J. C., Wei, F., Pasternak, S., Liang, C., Zhang, J., Fulton, L., Graves, T. A., Minx, P., Reily, A. D., Courtney, L., Kruchowski, S. S., Tomlinson, C., Strong, C., Delehaunty, K., Fronick, C., Courtney, B., Wilson, R. K. (2009). The B73 Maize Genome: Complexity, Diversity, and Dynamics. Science, 326(5956), 1112–1115. doi:10.1126/science.1178534

Stanke, M., Schöffmann, O., Morgenstern, B., & Waack, S. (2006). Gene prediction in eukaryotes with a generalized hidden Markov model that uses hints from external sources. BMC Bioinformatics, 7(1), 62. 10.1186/1471-2105-7-62

Verhoeven, E. C., & Anderson, N. P. (2020). Seed Production Research.

Wang, L.-G., Lam, T. T.-Y., Xu, S., Dai, Z., Zhou, L., Feng, T., Guo, P., Dunn, C. W., Jones, B. R., Bradley, T., Zhu, H., Guan, Y., Jiang, Y., & Yu, G. (2019). Treeio: An R Package for Phylogenetic Tree Input and Output with Richly Annotated and Associated Data. Molecular Biology and Evolution, 37(2), 599–603. 10.1093/molbev/msz240

Xu, S., Li, L., Luo, X., Chen, M., Tang, W., Zhan, L., Dai, Z., Lam, T. T., Guan, Y., & Yu, G. Ggtree: A serialized data object for visualization of a phylogenetic tree and annotation data. iMeta, n/a(n/a), e56. 10.1002/imt2.56

Zhou, C., McCarthy, S. A., & Durbin, R. (2022). YaHS: yet another Hi-C scaffolding tool. bioRxiv, 2022.2006.2009.495093. 10.1101/2022.06.09.495093

